# Mitophagosomes induced during EV-D68 infection promote viral nonlytic release

**DOI:** 10.1101/2024.12.05.627125

**Authors:** Alagie Jassey, Bimal Paudel, Michael A. Wagner, Noah Pollack, I-Ting Cheng, Raquel Godoy-Ruiz, David J. Weber, William T. Jackson

**Affiliations:** Department of Microbiology and Immunology and Center for Pathogen Research, University of Maryland School of Medicine, 685 W. Baltimore Avenue, Baltimore, MD 21201, USA; Department of Biochemistry and Molecular Biology, University of Maryland School of Medicine, Baltimore; Institute for Bioscience and Biotechnology Research, Rockville; The Center for Biomolecular Therapeutics, University of Maryland School of Medicine, Baltimore

**Author notes:** These authors contributed equally (Alagie Jassey, Bimal Paudel).

**Keywords:** Mitophagy, Parkin, PINK1, BCL2L13, Mitofusin-2, EV-D68

## Abstract

Enterovirus-D68 (EV-D68) is a plus-strand RNA virus that primarily causes infant respiratory infections. In rare pediatric cases, infection with EV-D68 has been associated with acute flaccid myelitis, a polio-like paralytic disease. We have previously demonstrated that EV-D68 induces nonselective autophagy for its benefit. Here, we demonstrate that EV-D68 induces mitophagy, the specific autophagic degradation of mitochondria. EV-D68 infection induces mitophagosome formation and several hallmarks of mitophagy, including mitochondrial fragmentation, mitochondrial membrane potential loss, and Parkin translocation to the mitochondria were observed in EV-D68 infected cells. The 3C protease of EV-D68 cleaves the mitochondrial fusion protein, mitofusin-2, near the C-terminal HR2 domain to induce mitochondrial fragmentation, and these fragmented mitochondria colocalized with double-stranded RNA (dsRNA), which labels viral RNA replication sites after peak viral RNA replication. Depleting components of mitophagy signaling specifically reduced EV-D68 release without impacting viral intracellular titers. Our results suggest that whereas the machinery of macroautophagy supports various stages of enterovirus replication, including viral genomic RNA replication and capsid maturation, mitophagy is the specific form of autophagy that regulates the nonlytic release of enteroviruses from cells.

**Significance:** EV-D68 is a respiratory pathogen and a significant cause of acute respiratory illness in infants, against which no prophylactic vaccines or antivirals exist. We have previously shown that EV-D68, like other enteroviruses, triggers and benefits from nonselective autophagy. However, it was unclear whether the virus induces selective autophagy. Here, we demonstrate that EV-D68 infection induces mitophagy and uses mitophagosomes to traffic virions to the surface of cells for nonlytic release. Our findings suggest that mitophagy may constitute the specific form of autophagy that facilitates the autophagosome-mediated nonlytic release of enteroviruses from infected cells and that targeting mitophagy could be an effective strategy to impede EV-D68 nonlytic release.

## Introduction

Enterovirus-D68 (EV-D68), a member of the *Picornaviridae* family, was first identified in 1962 in California [1]. Since its discovery, the virus has caused seasonal clusters of respiratory illnesses, particularly in children, and cases have risen in recent decades [2]. In the past decade, infection with EV-D68 has been associated with Acute Flaccid Myelitis (AFM), a polio-like neurological illness, that causes asymmetric paralysis in infants and is of significant public health concern [3,4]. To date, there is no prophylactic vaccine or antivirals against EV-D68 infection, and therefore, dissecting the mechanisms by which EV-D68 hijacks cellular processes/pathways to aid its reproduction could open opportunities for novel therapeutic interventions against the picornavirus.

EV-D68 is a positive sense single-stranded RNA virus with a genome size of 7.5 kb. The genome is encoded as a single polyprotein, which is processed by viral proteases to structural and nonstructural proteins. The structural proteins encapsidate the viral genome and play essential roles in viral binding and entry into the host cells. The nonstructural proteins play various roles during infection, including rearranging cellular membranes to form viral replication complexes. Picornavirus replication occurs on single or double-membrane vesicles [5]. These vesicles resemble autophagosomes, and many studies suggest that picornaviruses use autophagosomes for RNA replication, maturation, and nonlytic release [6–8]. However, the specific type or types of autophagy elicited during most picornavirus infections, including EV-D68, are unclear.

Mitochondria are dynamic organelles that undergo fusion and fission to facilitate mitochondrial redistribution and maintain a healthy mitochondrial network [9,10]. While mitochondrial fusion, orchestrated for the most part by mitofusin (MFN)-1 and −2, is known to inhibit mitophagy, mitochondrial fission, which is facilitated by Dynamin-related protein-1 (Drp-1), among others, promotes mitophagy [11]. Mitophagy is a specific type of autophagy that targets damaged and dysfunctional mitochondria for degradation, thereby maintaining mitochondrial quality control [12]. Mitophagy, in many ways, is similar to autophagy as they both involve the formation of the autophagosome, which engulfs the cargo and targets them to lysosomal for degradation. However, unlike nonselective autophagy, mitophagy involves additional molecular players, including Parkin, an E3 ubiquitin ligase, and PINK1, a mitochondrial health sensor, neither of which are implicated in nonselective bulk autophagy. In polarized mitochondria, PINK1 is constitutively imported to the inner mitochondrial membrane but it is rapidly degraded by the mitochondrial protease PARL [13]. However, in depolarized mitochondria, PINK1 import to the inner mitochondrial membrane is inhibited, leading to its accumulation in the outer mitochondrial membrane and recruitment of Parkin to the mitochondria. Parkin then ubiquitinates the mitochondria, leading to the autophagic degradation of the mitochondria [14,15]. Parkin translocation to mitochondria is a hallmark of Parkin/PINK1-dependent mitophagy and is often used to determine mitophagy induction in cell culture.

Besides Parkin/PINK1-dependent mitophagy, other forms of mitophagy, orchestrated by specific mitophagy receptors, exist [16]. For instance, recently, Bcl-2-like protein 13 (BCL2L13), the mammalian homologue of yeast ATG32, which is vital for mitophagy induction in yeast, was identified as a mitophagy receptor in mammalian cells [17]. BCL2L13 binds directly to LC3 through its LC3 interacting region and induces mitochondrial fragmentation and mitophagy in Drp1- and Parkin-depleted cells, indicating that mitophagy induced by BCL2l13 does not require Drp1 and Parkin.

Here, we demonstrate for the first time that EV-D68 infection induces mitophagosome formation, and these mitophagosomes specifically promote EV-D68 nonlytic release from cells. Knockdown of components of mitophagy (both receptor-mediated and Parkin/PINK1-dependent mitophagy) signaling decreased viral release without significantly affecting intracellular viral titers. Our results suggest that mitophagosomes may be the specific form of autophagosomes responsible for the nonlytic release of enteroviruses.

## Results

### EV-D68 depolarizes mitochondria and induces Parkin translocation to mitochondria

EV-D68, like several other picornaviruses, is known to induce and hijack the autophagic machinery for its benefit. However, whether or not EV-D68 induces specific organellar autophagy is unclear. CVB3, a related picornavirus and a significant cause of myocarditis, is known to induce mitophagy and uses mitophagosomes for viral replication and egress [18]. Since mitophagy is typically preceded by mitochondrial damage/depolarization, we asked whether EV-D68 infection affects mitochondrial dynamics. For this purpose, we mock-infected or infected H1HeLa cells and measured the mitochondrial membrane potential. We included Carbonyl cyanide m-chlorophenyl hydrazone (CCCP), a drug used widely to induce mitophagy as a positive control. As shown in Fig. 1A, EV-D68 causes mitochondrial depolarization at similar levels as the CCCP control. To understand how EV-D68 causes mitochondrial depolarization, we examine the effect of EV-D68 infection on intracellular calcium levels. An increase in intracellular calcium levels increases mitochondrial calcium uptake, leading to mitochondrial membrane potential loss. We observed an increase in intracellular calcium levels in EV-D68-infected and CCCP-treated cells compared to the control, suggesting that an increase in intracellular calcium levels may contribute to mitochondrial depolarization during EV-D68 infection (Fig. 1B). Next, we examined the effect of EV-D68 infection on the morphology of the mitochondrial network by immunofluorescence assay (IFA). In contrast to the mock control, which shows a normal tubular mitochondrial network, as expected, we observed extensive mitochondrial network fragmentation in EV-D68-infected cells (Fig. 1C). We then examined the effect of EV-D68 infection on Parkin translocation to the mitochondria, a hallmark of mitophagy. While there were no significant changes in Parkin levels between the mock-infected and EV-D68-infected cells in the non-mitochondrial fraction, EV-D68 infection increases Parkin translocation to the mitochondria to a similar degree as the CCCP control (Fig. 1D).

**Figure 1.**
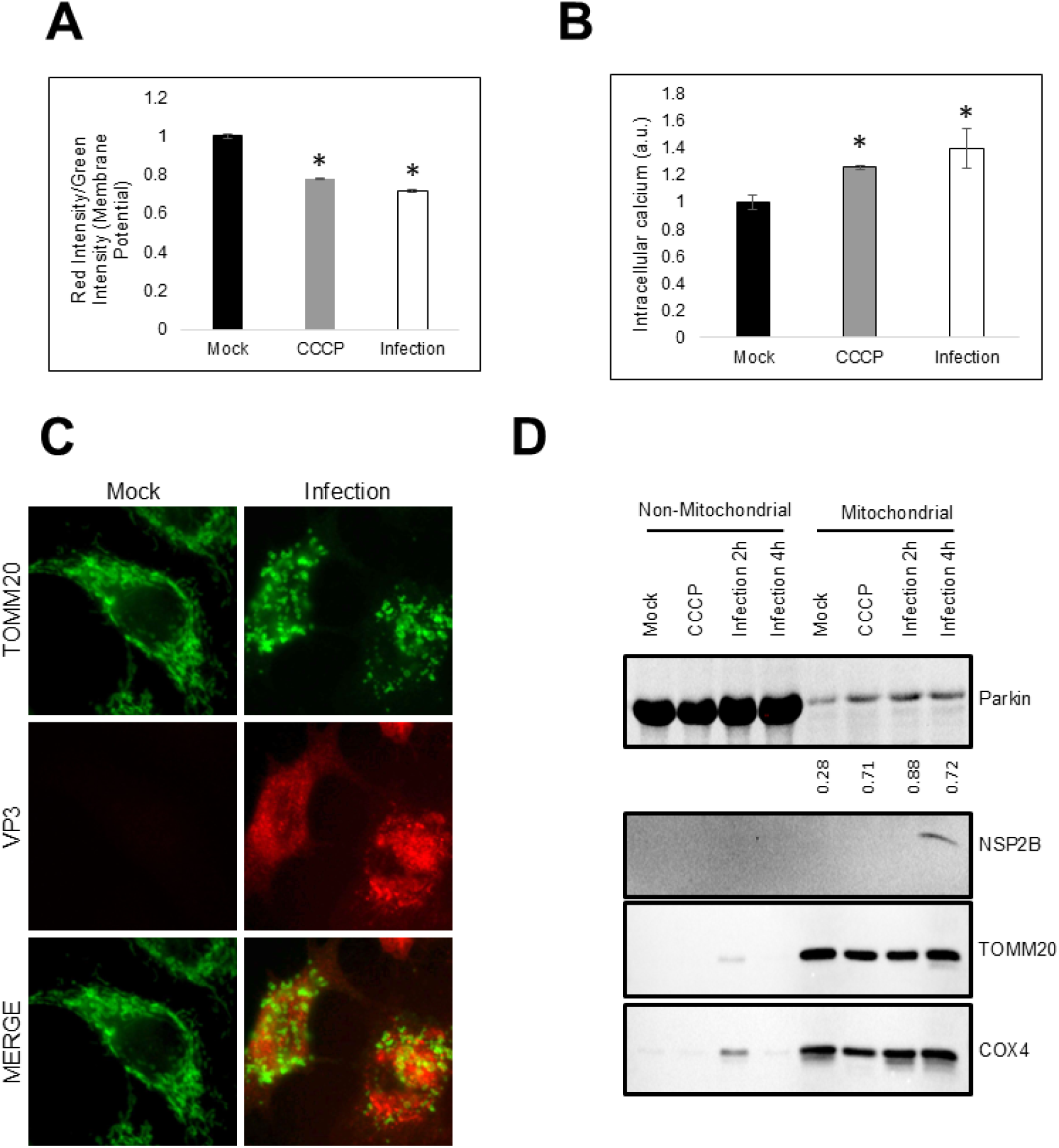
Enterovirus D68 depolarizes mitochondria and induces Parkin translocation to the mitochondria. (A) H1HeLa cells were either treated with 10 µM CCCP (4h) or infected with EV-D68 at an MOI of 25 for 4 h. Mitochondrial membrane potential was determined using JC10 mitochondrial membrane potential kit. (B) H1HeLa cells were treated with CCCP for 12 h or infected with EVD-68 for 4 h and the intracellular calcium levels were measured by Fluo-8 Calcium Assay Kit – No Wash. (C) A549 cells were either mock-infected or infected with EV-D68 (MOI= 25) for 5 h. The cells were then fixed and IFA was performed against TOMM20. (D) H1HeLa cells were treated with CCCP or infected with EV-D68 (MOI =25) for the indicated time points. The cells were then fractionated using a mitochondrial fractionation kit and a western blot was performed against the indicated antibodies. Error bars indicate mean ± SEM of 3 independent repeats. Unpaired student’s t-test was used for the statistical analyses (* = p ≤ 0.05.).

### EV-D68 induces mitophagosome formation

Another hallmark of mitophagy is the detection of mitophagosomes (mitochondria enveloped in autophagosomes) by electron microscopy. To further confirm mitophagy induction during EV-d68 infection, we mock-infected or infected H1HeLa cells with EV-D68 (MOI =25) for 4h for electron microscopy. As shown in Fig. 2A, unlike the mock infection control, which shows normal cell morphology and organelle structures, EV-D68 rearranges membranes, resulting in the so-called replication organelle formation, characteristic of picornavirus infection. Zooming in on the EM images, we observed mitochondria with intact cristae in the mock-infected control, in contrast to EV-D68 infected cells, in which loss of mitochondrial cristae was evident (Fig. 2B). Furthermore, we observed mitophagosomes in EV-D68 infected cells (Fig. 2C). Together, these data suggest that EV-D68 disrupts mitochondrial dynamics and induces mitophagosome formation.

**Figure 2.**
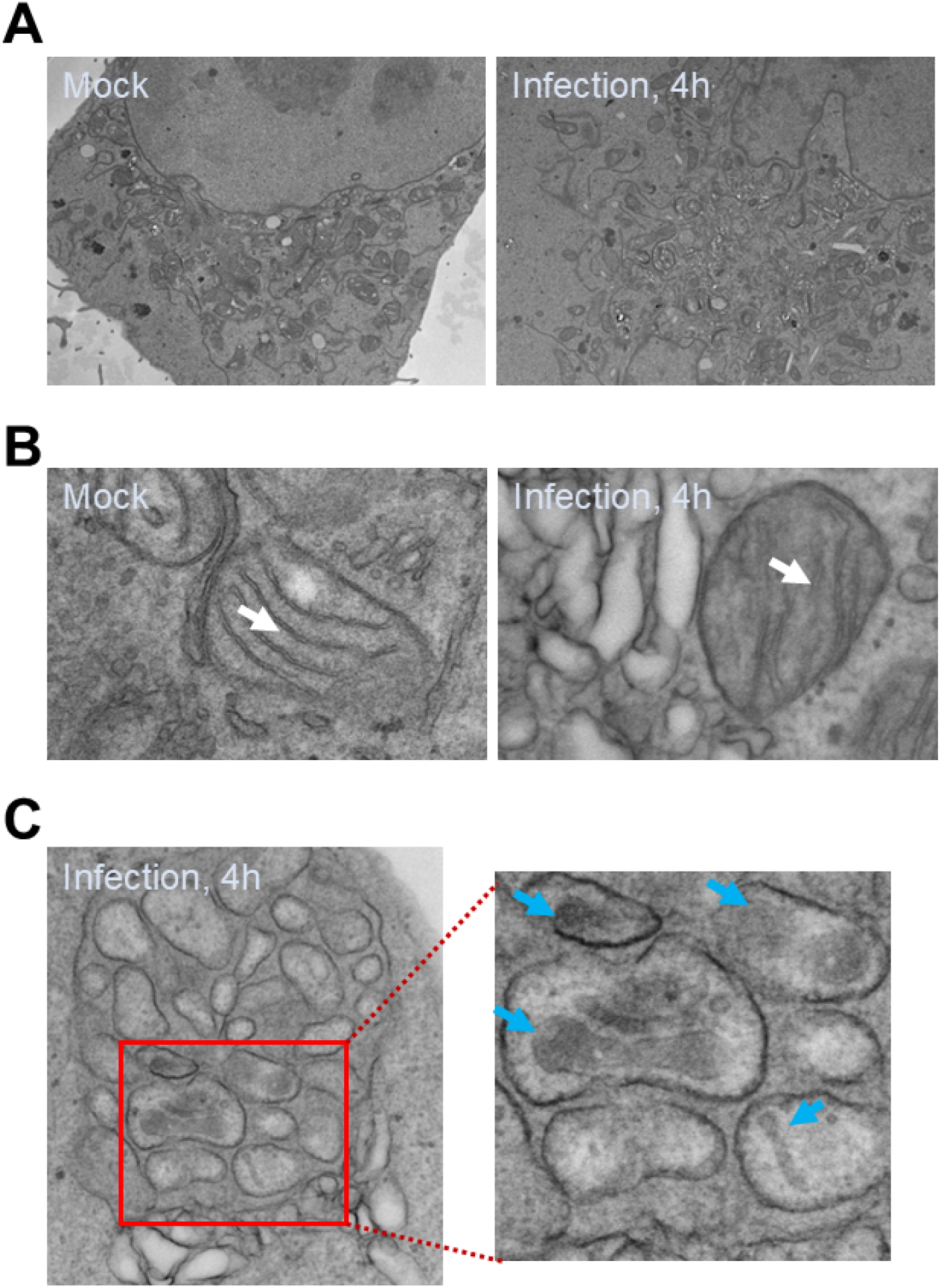
EV-D68 infection induces mitophagosome formation. H1HeLa cells were mock-infected or infected with EV-D68 (MOI =30) for 4 h for EM imaging. (A) Normal cell morphology and intact mitochondrial network in mock-infected cells (left) vs membrane rearrangement and replication organelle formation in EV-D68 infected cells (right). (B) Loss of mitochondrial cristae in EV-D68-infected cells compared to the mock cells, shown with white arrows. (C) Mitophagosomes in EVD-68 infected cells are shown with blue arrows. Scale bar = 2 µm.

### Mitochondria colocalize to viral RNA replication sites late in EV-D68 infection

The results in Fig. 1C demonstrate that EV-D68 infection causes a fragmentation of the mitochondrial network. We wanted to understand the timing of mitochondrial fragmentation during infection and whether the mitochondrial membranes and mitophagosomes are used as sites of viral RNA replication. We mock-infected or infected cells with EV-D68 and performed an IFA against translocase of outer mitochondrial membrane 20 (TOMM20), an outer mitochondrial membrane protein and double-stranded RNA (dsRNA), a viral RNA replication intermediate commonly used to mark viral RNA replication sites. As indicated in Fig. 3, mock-infected cells show a normal tubular mitochondrial signal. In contrast, EV-D68 infection causes fragmentation of the mitochondrial network, which starts early in the infection (∼3hpi) and becomes more apparent as infection progresses (∼5hpi). While there was no significant colocalization between the dsRNA signals and mitochondria at 3hpi, we observed an increase in colocalization between these signals at 5hpi.

**Figure 3.**
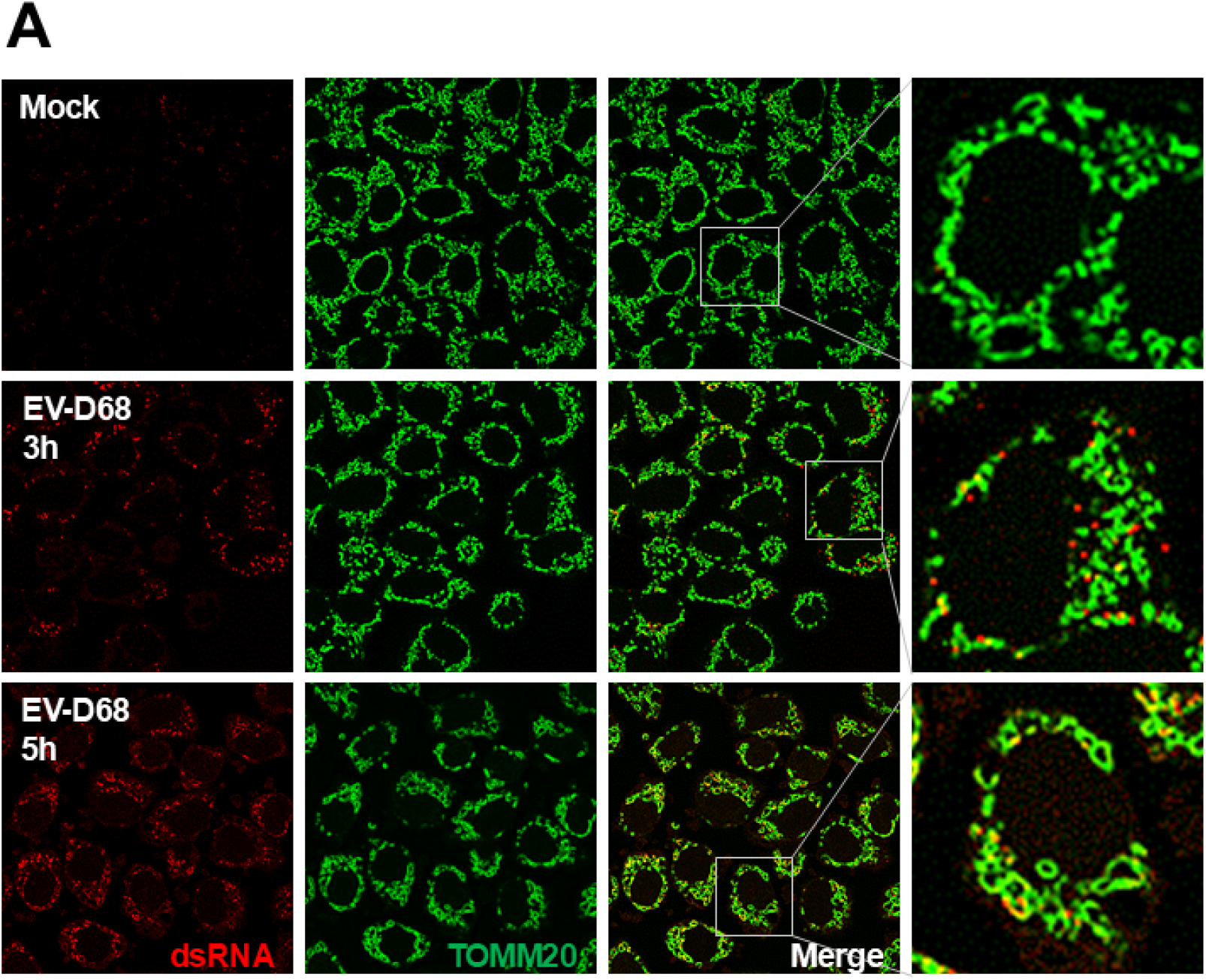
Mitochondria colocalizes to viral dsRNA late during EV-D68 infection. H1HeLa cells were infected with EV-D68 (MOI =25) for the indicated time points. The cells were fixed and subjected to IFA against dsRNA and TOMM20.

### Enterovirus D68 cleaves the mitochondrial fusion protein, mitofusin-2 (Mfn-2) at C-terminal HR2 domain

CVB3 has previously been shown to induce mitochondrial fragmentation and mitophagy by inducing dynamin-related protein-1 (Drp-1) phosphorylation. To understand how EV-D68 induces mitochondrial fragmentation and mitophagy, we examined the expression levels of proteins that regulate mitochondrial dynamics, including MFN-2 and Drp-1, upon EV-D68 infection. While EV-D68 infection does not significantly alter Drp-1 levels, we observed, in addition to the full-length MFN-2 band, a smaller band beginning at 4hpi, suggesting a potential cleavage fragment (Fig. 4A and Fig. S1A). In contrast, we did not detect any potential cleavage fragments for MFN-1 (Fig. 4A). The picornaviruses 3C-protease is known to cleave several host proteins, including the autophagosomal SNARE protein, SNAP29. To test if the 3C protease is responsible for MFN-2 cleavage during EV-D68 infection, we incubated cell lysate (with and without MFN-2 overexpression) with EV-D68-3C-protease in vitro. Our in vitro cleavage assay shows that 3C-protease cleaves MFN-2, displaying a similar band pattern as in the viral infection (Fig. 4C). Based on the size of the cleavage fragment, we hypothesize that the cleavage may occur closer to the C-terminal end, which contains the Heptad repeat 2 (HR2) domain essential for mitochondrial dimerization. To confirm this hypothesis, we overexpressed a C-terminally HA-Tagged MFN-2 plasmid in H1HeLa cells and left the cells uninfected or infected with EV-D68 for western blot against HA and MFN-2. As expected, we detected the 65 kDa cleavage fragment with an anti-MFN-2 antibody (Fig. 4D). In contrast, the anti-HA antibody failed to detect the 65 kDa cleavage fragment but instead stained a smaller 17 kDa fragment (Fig. 4E), indicating that EV-D68 3C protease cleaves MFN-2 at the C-terminus. We wanted to confirm whether MFN-2 cleavage is conserved among enteroviruses. We infected cells with different enteroviruses and included EV-D68 as a control. As shown in Fig. 4B, all the enteroviruses tested cleaved MFN-2 at similar levels, suggesting that mitofusin-2 cleavage may be conserved amongst enteroviruses.

**Figure 4.**
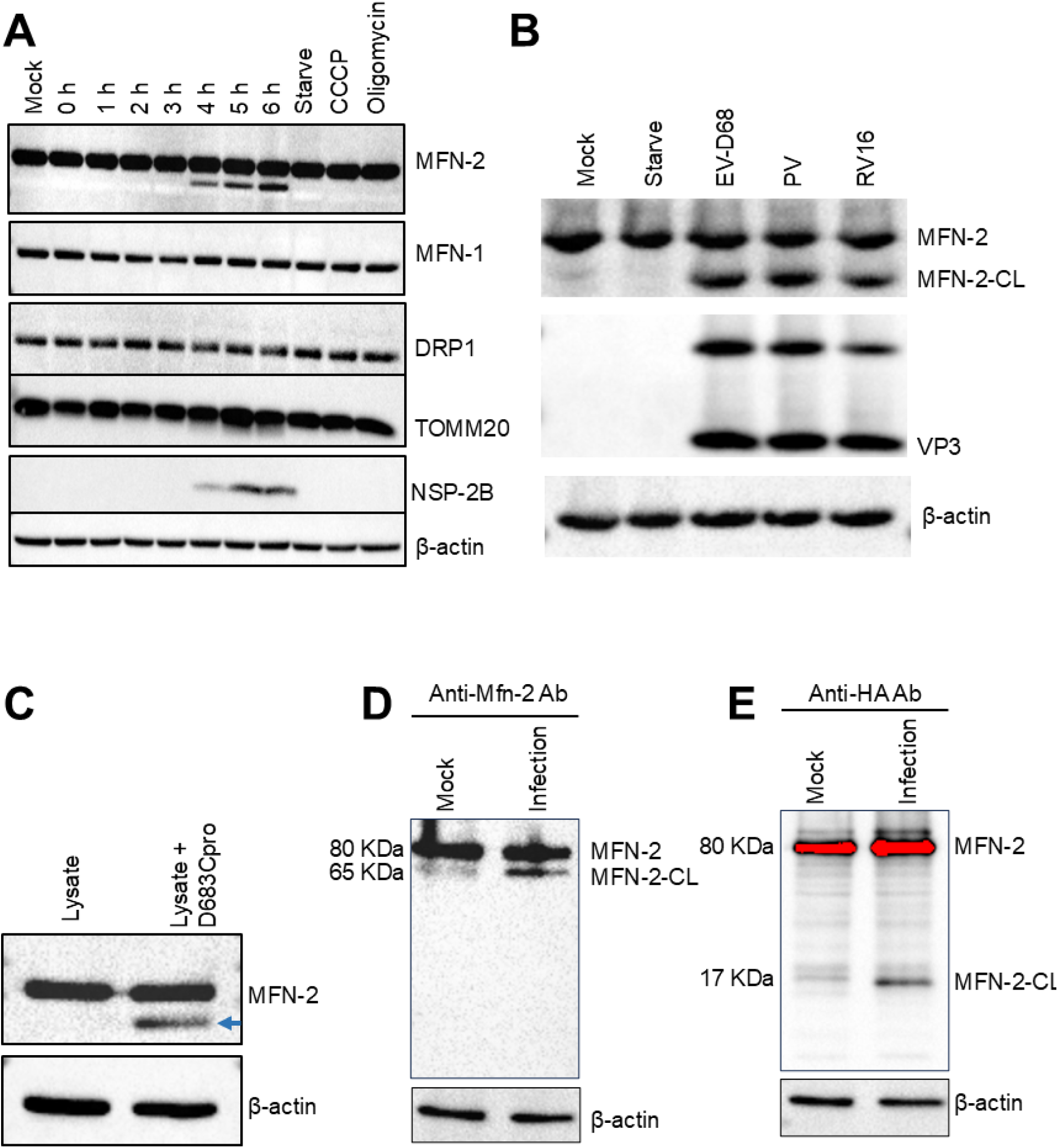
EV-D68 cleaves the mitochondrial fusion protein, MFN-2. (A) H1HeLa cells were infected for the indicated time points with EV-D68 (MOI =25). Western blot was then performed against the mitochondria-associated proteins. Lysates from cells treated with CCCP or Oligomycin-A for 12h were included as positive controls. (B) H1HeLa cells were infected with the indicated enteroviruses for 5 h, after which lysates were prepared for western blot. (C) A549 cell lysates were incubated with and without the 3C protease for an in vitro cleavage assay for western blot against MFN-2. (D and E) H1HeLa cells were transfected with pcDNA3.1 MFN-2-HA for 24 h. The cells were either mock-infected or infected with EV-D68 (MOI =25) for 5 h for western blot using anti-MFN-2 (4D) or anti-HA (4E) antibodies.

### Active mitophagy signaling is essential for EV-D68 release

The finding that the dsRNA signal colocalizes with TOMM20 at 5hpi, but not 3hpi (Fig. 3), suggests that mitophagy may be more important for the later stages of the viral life cycle post-RNA replication. To test this hypothesis, we knocked down Parkin in H1HeLa cells, infected the cells and performed a plaque assay to measure intracellular and extracellular viruses. Knockdown of Parkin had no significant effect on viral intracellular titers but markedly reduced extracellular titers (Fig. 5A). We observed similar results in BCL2L13 knockdown H1HeLa cells, wherein knockdown of the mitophagy receptor reduced extracellular viral titers, without affecting viral intracellular titers (Fig. 5B). We also knockdown Parkin, PINK1 (Fig. 5C) and BCL2L13 (Fig. 5D) in A549 cells and observed similar results. Next, we wanted to test whether simultaneous knockdown of Parkin/PINK1 mitophagy and receptor-mediated mitophagy through BCL2L13 knockdown will have additive effects on the viral titers. As shown in Fig. 5E, simultaneous depletion of BCL2L13 and Parkin or BCL2L13 and PINK1 did not further reduce EV-D68 titers compared to the individual knockdowns in A549 cells. Together, these results indicate that mitophagy specifically regulates EV-D68 release.

**Figure 5.**
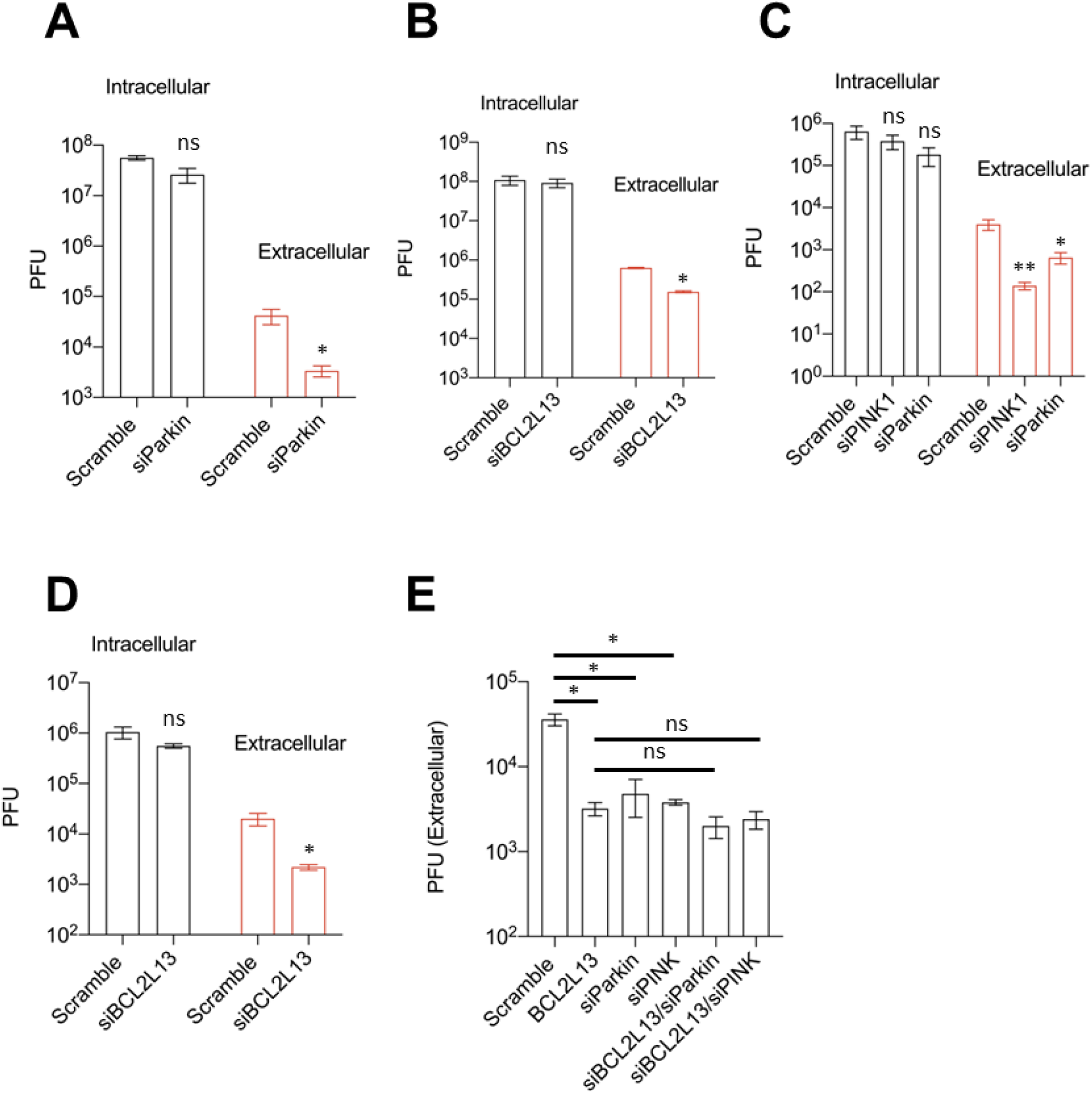
Active mitophagy signaling is essential for EV-D68 release. (A) H1HeLa cells were transfected with the scramble and Parkin siRNAs for 48 h. The cells were then infected with EV-D68 for 5 h and viral titers were measured by a plaque assay. (B) H1HeLa cells were transfected with siRNAs against the mitophagy receptor, BCL2L13 for 48 h. The cells were then infected as in A for a plaque assay-based viral titer measurement. (C) A549 cells were transfected with scramble, Parkin, and PINK1 siRNAs for 48 h. The cells were infected with EV-D68 for 5h and viral titers were determined by a plaque assay. (D) A549 cells were transfected with the indicated siRNAs for 48 h. The cells were then infected and tittered as in A. (E) A549 cells were transfected with the indicated mitophagy-associated siRNAs for 48 h, followed by viral infection and titer determination as in A. An MOI of 0.1 was used for all infections. Error bars indicate mean ± SEM of at least 2 independent repeats. Unpaired student’s t-test was used for the statistical analyses (**= p< 0.01; *= p< 0.05; ns=not significant.).

### Knockdown or overexpression of mitofusin-2 attenuates EV-D68 release

Our data shows that MFN-2 is cleaved late during EV-D68 infection by the viral 3C protease, suggesting that mitofusin-2 may be detrimental to EV-D68 release. We depleted MFN-2 in H1HeLa and A549 cells and infected them with EV-D68 for viral titer measurement. To our surprise, the depletion of MFN-2, much like the knockdown of Parkin and PINK1, reduced EV-D68 extracellular titers in both H1HeLa (Fig. 6A) and A549 cells (Fig. 6B), without significantly impacting viral intracellular titers. To uncover how MFN-2 depletion attenuates EV-D68 release, we knocked down MFN-2 in H1HeLa and A549 cells for a western blot against TOMM20, an outer mitochondrial membrane protein, whose degradation is often used as an indication of active mitophagy. As demonstrated in Fig. 6C, the knockdown of mitofusin-2 leads to a loss of TOMM20 staining in both H1HeLa and A549 cells. In contrast, EV-D68 infection of these cells does not reduce TOMM20 staining (Fig. 6D). These findings suggest that the depletion of TOMM20 causes mitochondrial degradation, depleting the mitophagosomes necessary for viral egress and consequently restricting viral release. To further understand how mitofusin-2 regulates EV-D68 infection, we overexpress mitofusin-2 in H1HeLa cells (Fig. 6E) followed by viral infection (Fig. 6F). Intriguingly, overexpression of mitofusin-2 also reduced viral extracellular titers and had major effects on intracellular viral titers (Fig. 6F).

**Figure 6.**
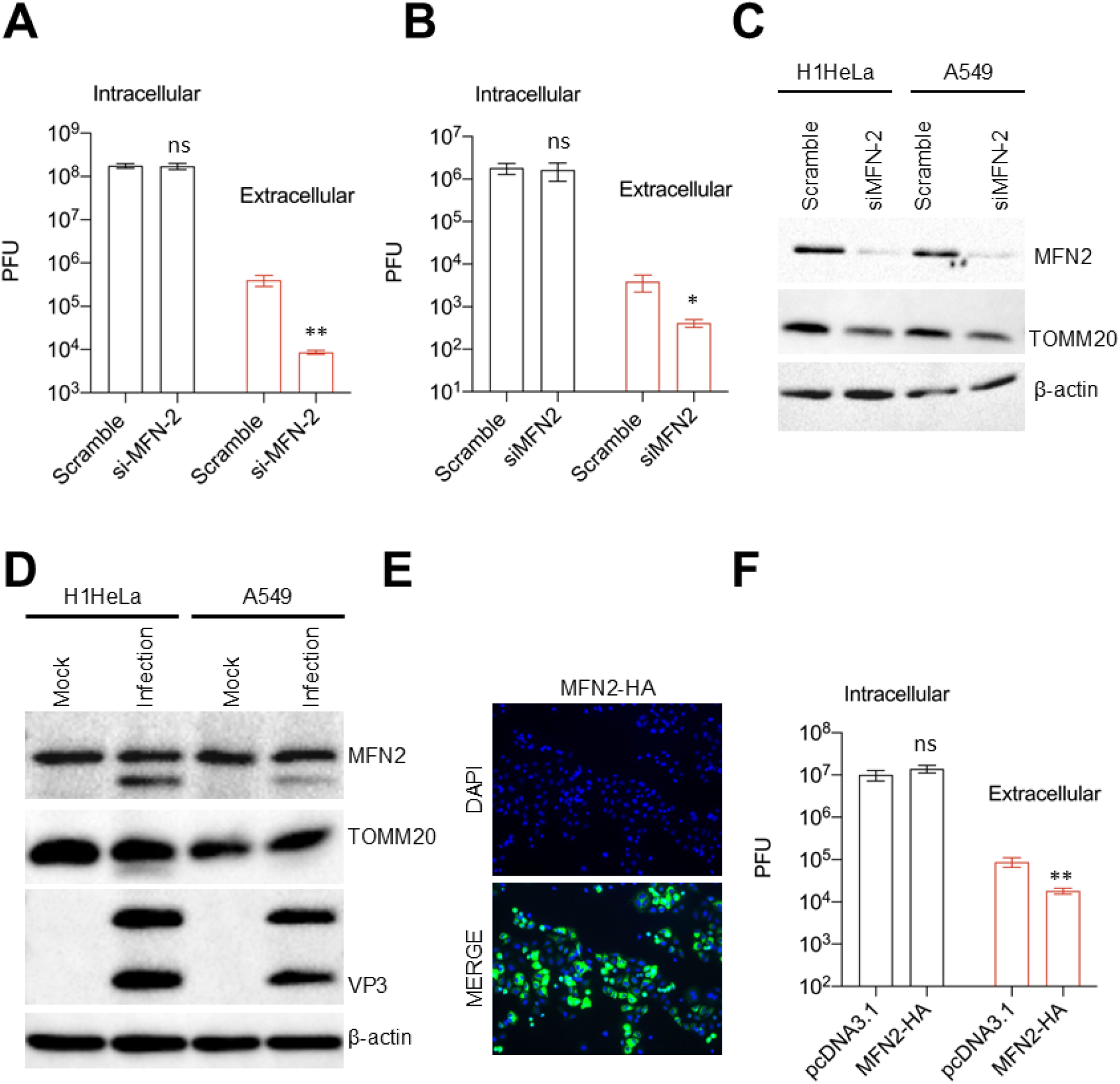
Knockdown or overexpression of mitofusin-2 attenuated EV-D68 release. (A) H1HeLa cells were transfected with scramble control or mitofusin-2 siRNAs for 48 h. The cells were then infected with EV-D68 for 5 h, followed by viral titer measurement by a plaque assay. (B) A549 cells were transfected and infected as in A for viral titer determination by a plaque assay. (C) H1HeLa and A549 cells were transfected with the indicated siRNAs for 48 h, after which the whole cell lysates were collected and prepared for a western blot against the indicated antibodies. (D) H1HeLa and A549 cells were infected with EV-D68 (MOI =30) for 5 h for western blot against mitofusin-2 and VP3. (E and F) H1HeLa cells were transfected with either a pcDNA3.1 control plasmid or MFN2-HA plasmid for 24 h (E). The cells were then infected with EV-D68 (MOI = 0.1) for 5 h (F) and viral titers were determined by a plaque assay. Error bars indicate mean ± SEM of at least 2 independent repeats. Unpaired student’s t-test was used for the statistical analyses (**= p< 0.01; *= p< 0.05; ns=not significant.).

## Discussion

Although we have previously demonstrated that EV-D68 subverts autophagy for its replication, whether the virus induces organelle-specific autophagy was unclear. We show here for the first time that EV-D68 infection induces mitophagy. However, unlike mitophagy induced by MFN-2 depletion, which causes complete mitophagy as evidenced by the reduction in TOMM20 staining in MFN-2-depleted cells (Fig. 6C), EV-D68 infection induces incomplete mitophagy and uses the mitophagosomes for nonlytic viral egress. Our data suggests that mitophagosomes constitute the specific form of autophagosomes that regulate the autophagosome-mediated exit without lysis of enteroviruses.

Our EM data shows a loss of mitochondrial cristae, the main site of eukaryotes’ biological energy conversion, in EV-D68-infected cells, suggesting that EV-D68 infection causes mitochondrial damage/dysfunction (Fig. 2B) [19]. Consistent with mitochondrial damage/dysfunction, we found that EV-D68 infection causes mitochondrial membrane potential loss and mitochondrial fragmentation (Fig. 1A, 1C, and Fig. 3). As damaged mitochondria are significant sources of reactive oxygen species, which are harmful to macromolecules, their timely removal by mitophagy could maintain mitochondrial health and cell viability [20,21]. Given that EV-D68, similar to many viruses, is adapted to replicating in viable cells, the induction of mitophagy during EV-D68 infection could enhance genomic RNA replication.

Although our data demonstrate that EV-D68 infection causes mitochondrial damage, how EV-D68 damages mitochondria and the specific viral factors involved are unclear. Hepatitis C Virus (HCV), a member of the Flaviviridae family and an important cause of end-stage liver disease, was previously shown to cause mitochondrial damage by inducing ROS production. The viral multifunctional structural (Core) and nonstructural (NS5A) proteins were identified as the essential inducers of ROS during HCV infection [22–24]. Transient overexpression of HCV NS5A induces mitochondrial fragmentation, and inhibition of ROS attenuated the NS5A-induced mitophagy, suggesting a possible link between ROS generation and mitochondrial fragmentation [25,26]. Interestingly, the 2B protein of EV-A71 was reported to trigger ROS production by interacting with the outer mitochondrial membrane protein, voltage-dependent anion channel 3 (VDAC3), which is essential for viral RNA replication [27]. We also detected EV-D68 2B protein in the mitochondrial fraction (Fig. 1D), suggesting a possible interaction between the viral protein and mitochondria. It will be interesting to uncover if the 2B protein of EV-D68 is involved in ROS generation and EV-D68-induced mitochondrial damage and mitophagy.

HCV was also shown to cause mitochondrial fragmentation and mitophagy to dampen the type I interferon response by inducing Drp-1 protein expression and phosphorylation [28]. CVB3 similarly induces Drp-1-dependent mitochondrial fragmentation and mitophagy. However, unlike HCV, CVB3 uses mitophagosomes for release, akin to the autophagosome-media exit without lysis model during poliovirus infection [8,18]. Unlike these viruses, the EV-D68-mediated mitochondrial fragmentation does not appear to depend on Drp-1, as we did not observe any significant change in Drp-1 protein levels during infection. Instead, our findings suggest that EV-D68 induces mitochondrial fragmentation by cleaving MFN-2, a vital mitochondrial fusion protein (Fig. 4). MFN-2 cleavage begins around 4hpi during EV-D68 infection, which, interestingly, is the earliest time point we observed appreciable mitochondrial network fragmentation (Fig. 4A). Through in vitro cleavage assay, we show that the viral 3C protease cleaves MFN-2 near the C-terminal HR2 domain, essential for tethering MFN from opposing outer mitochondrial membranes, mediating mitochondrial fusion, separating the HR2 domain from the GTPase domain at the N-terminus, which also important mitochondrial fusion [29]. Therefore, by cleaving MFN-2 at or near the HR2 domain, EV-D68 effectively blocks mitochondrial fusion, leading to mitochondrial network fragmentation.

In contrast to bulk nonselective autophagy, which is known to promote various stages of the enterovirus life cycle, including RNA replication, our results show that mitophagy is not crucial for EV-D68 RNA replication. Knockdown of Parkin/PINK1- and receptor-mediated mitophagy did not affect EV-D68 intracellular titers (Fig. 5A, B, C, and D). Instead, knockdown of the cellular mitochondrial quality control process specifically impaired EV-D68 nonlytic release, suggesting that the mitophagosomes generated during EV-D68 infection are used to transport virions out of cells in a nonlytic fashion.

Unexpectedly, we also found that depleting MFN-2 phenocopies mitophagy knockdown, reducing viral release without impacting intracellular viral titers (Fig. 6A and B). This was surprising because we reasoned that the knockdown of MFN-2, known to induce fragmentation of mitochondrial networks, will induce mitophagosomes and promote EV-D68 release. However, as shown in Fig. 6C, the knockdown of MFN-2 leads to degradative mitophagy indicated by the loss of TOMM20 staining in MFN-2-depleted cells. This finding suggests that by allowing mitophagy flux induction before the initiation of viral infection, MFN-2 knockdown deprives the virus of these crucial vesicles and consequently restricts EV-D68 release.

Equally unexpected was our observation that overexpression of MFN-2 also reduces EV-D68 release (Fig. 6F). Precisely how MFN-2 overexpression attenuates EV-D68 release is unclear. However, in light of the finding that mitochondrial ER coupling orchestrated by MFN-2 inhibits mitophagy initiation[30], we posit that overexpression of MFN-2 will enhance ER-mitochondria tethering and prevent mitophagosome formation during EV-D68 infection, thereby reducing viral release.

## Materials and Methods

### Cell culture

H1HeLa (ATCC, CRL-1958), A549 (ATCC, CCL-185) were cultured in DMEM (Gibco, 11965-092) supplemented with 10% heat-inactivated fetal bovine serum albumin (GemininBio, 100-106), 1x penicillin-streptomycin (Gibco, 10, 378-016), and 1x sodium pyruvate (Gibco, 11, 360-070). Cultured cells were incubated in a 5% CO2 incubator at 37°C.

### Plasmids

The pcDNA3.1 MFN2-HA plasmid (plasmid #139192) was obtained from Addgene (plasmid #139192). Transfection of the plasmids into the desired cells was done by using the Lipofectamine 2000 transfection reagent (Invitrogen, 11, 668-019). The transfection complex was removed at 6 h post-transfection and replaced with basal DMEM media. Follow-up treatments/infection was initiated 24 h post DNA transfection.

### Western blotting

Cells were grown in 6-well plates and collected in PBS (Quality Biological, 114 −058-101), after treatment/infection. Collected cells in microcentrifuge tubes were lysed using RIPA buffer (Sigma, R0278) supplemented with Complete Tablets Mini Protease Inhibitor Cocktail (ROCHE, 11, 836, 170, 001) and incubated on ice for 30 minutes. Protein lysates were collected after centrifuging the tubes at 10,625 x g for 30 minutes. Protein concentration was measured using Bradford assay, and the lysates were then loaded to SDS-PAGE. After SDS-PAGE, proteins were transferred to polyvinylidene fluoride (PVDF; Bio-Rad, 1, 620, 177) membranes. The membranes were then blocked for 1 h with 5% skim milk, washed twice (10 minutes each) with TBST (100 mM Tris-hydrochloride, pH 7.4, 2.5 M sodium chloride, and 0.125% Tween-20 [Sigma Aldrich, P1379]), and incubated with primary antibodies overnight on a shaker at 4°C. The membranes were washed twice with TBST and incubated with secondary antibodies (1:2000) at room temperature for 1 h before being developed by Western Lightning ECL. Images were acquired using Chemi-Doc (Bio-Rad).

### Immunofluorescence assay

Cells were grown in 12 well plates on glass coverslips. After the treatment, cells were fixed with 4% PFA. Cells were washed with 1x PBS and permeabilized using either 0.3% Triton-X or 1% Saponin, after which they were blocked for 1h at room temperature with 3% bovine serum albumin. Cells were incubated with primary antibodies overnight at 4°C. The next day, cells were washed 3 times (5 minutes per wash) with PBS and incubated with secondary antibodies for 1 h at room temperature, protected, from light. Coverslips with treated cells were mounted on glass slides with Prolong Glass Antifade Mountant with NucBlueTM, mounting media (Invitrogen, P36985). Prepared slides were imaged using Confocal or Revolve microscopes.

### Electron Microscopy

For electron microscopy analysis, cells were fixed with a solution of 2% paraformaldehyde, and 2.5% glutaraldehyde in 0.1 M PIPES buffer (pH 7.2), scrapped off the tissue culture vessel, washed in 0.1M PIPES buffer and collected by centrifugation. Cell pellets were enrobed in 2.5% low melting point agarose, trimmed into 1mm3 blocks and post-fixed with 1% osmium tetroxide and 1.5% potassium ferrocyanide in 0.1M PIPES buffer for 1 hour at 4°C. After washing, agarose blocks containing cells were en bloc stained with 1% uranyl acetate in water and dehydrated using increasing concentrations of ethanol from 30%; 50%; 70%; 90% and 100% for 10 min at each step. Specimens were then incubated with two changes of 100% acetone and infiltrated, in increasing concentration of Araldite-Epoxy resin (Araldite, EMbed 812; Electron Microscopy Sciences, PA), and embedded in pure resin at 60°C for 24 to 48 h. Ultrathin sections at ∼70nm thickness were cut using Leica UC6 ultramicrotome (Leica Microsystems, Inc., Bannockburn, IL), and examined in a FEI Tecnai T12 electron microscope operated at 80 kV. Digital images were acquired using a AMT bottom mount CCD camera and AMT600 software.

### Antibodies

The following primary antibodies were purchased and used in this study. Rabbit polyclonal-anti-Pink1 (Novus Biologicals, BC100-494). Rabbit polyclonal-anti-Parkin (Cell Signaling, #2132). Rabbit polyclonal-anti-BCL2l13 (Proteintech 16612-1-AP). Mouse monoclonal-anti-Sqstm1 (Abnova, H00008878-M01). Rabbit polyclonal-anti-LC3B (Novus Biologicals, NB600-1384). Rabbit monoclonal-anti-Mitofusin-2 (Cell Signaling, #9482). Rabbit monoclonal-anti-Mitofusin-1 (Cell Signaling, #14739). Rabbit monoclonal-anti-DRP1 (Cell Signaling, #5391). Rabbit monoclonal-anti-TOMM20 (Abcam, ab186735). Rabbit polyclonal-anti-Cox4 (Cell Signaling, #4844). Rabbit polyclonal-anti-EV-D68-NSP-2B (Invitrogen, PA5-112042). Mouse monoclonal-anti-ß-actin (Novus Biologicals, NB600-501). Mouse monoclonal-anti-dsRNA (Sigma Aldrich, MABE1134). HRP-conjugated anti-mouse secondary and anti-rabbit secondary antibodies were obtained from BioRad (#1706516 and #1706515 respectively).

### Virus infections

Viral infection was done following previously described methods with some modifications. An MOI of 0.1 (low) or 25 (high) with indicated time was used for all the infections. Briefly, virus particles were diluted in serum-free media, added to the cells, and incubated at 37° C for 30 minutes for adsorption. Serum-free media was removed, cells were washed with 1x PBS and basal media was added until the end of the infection. After the infection, the cells were either fixed and prepared for immunofluorescence assay or electron microscopy or collected in microcentrifuge tubes for western blot analysis or plaque assay for viral titer.

### siRNA and gene knockdowns

Parkin, PINK1, BCL2L13, and MFN-2 were knocked down using siRNAs. H1HeLa cells or A549 cells were grown to 40% confluency and transfected with siRNAs against the genes mentioned above or the scrambled siRNA (Mission siRNA universal negative control 1 [Sigma, SIC001]) using Lipofectamine 2000 and Opti-MEM (Gibco, 31,985-070). Briefly, for each well of 6-well plate, 100 to 300 nM of each siRNA and 5 uL of Lipofectamine (Invitrogen, 11,668-019) were separately incubated in Opti-MEM for 10 minutes, after which they were mixed and incubated for 1 h at room temperature. Then, the cells were washed with 1x PBS and incubated with the transfection mixture for 6 h, after which the transfection reagent was removed, and cells were washed with 1x PBS and supplemented with complete media. Follow-up experiments like virus infection or knockdown efficiency were performed after 48 h of transfection.

### Plaque assay

For virus titers, cells were scraped in 1x PBS using cell scrapers and transferred to microcentrifuge tubes followed by three freeze-thaw cycles. The resulting lysates were serially diluted in serum-free media and added to H1Hela cells for 30 minutes. After 30 minutes of adsorption, the inoculum was removed and overlaid with a 1:1 mixture of 2x MEM and 2% agar for 48 h, after which agar overlays were removed for crystal violet staining.

### Drugs and their treatments

Carbonyl cyanide 3-chlorophenylhydrazone (CCCP) (Sigma Aldrich, C2759) was used at a concentration of 10 µM for 4 h or 12 h (as indicated in the experiment). Oligomycin A (Sigma Aldrich, 75351) was used at 10 µM concentration for the indicated periods.

### RNA isolation and quantitative polymerase chain reaction (qPCR)

Total RNA was isolated from H1Hela or A549 cells using TRIZol (Ambion, 15,596,026) following the manufacturer’s instructions. Thermo Scientific RevertAid H Minus First Strand cDNA synthesis kit (K1632) was used to synthesize the cDNAs after the removal of genomic DNA. qPCR was done using KiCqStart SYBR qPCR Ready Mix (Sigma, KCQS01) with the 7500 Fast Dx Real-Time PCR Instrument (Applied Biosystems) according to the manufacturer’s protocols. Forward primer 5’ -TAACCCGTGTGTAGCTTGG-3’ and Reverse primer 5’-ATTAGCCGCATTCAGGGGC-3’, which are specific to 5’ untranslated region (UTR), were used to amplify the EV-D68 genome. The gene expression results were normalized to GAPDH using IDT Predesigned primers (Hs.PT.39a.22214836). Data was plotted as a relative expression of the viral genome compared to the 0 h infection time point, which is a data point collected after 30 minutes of virus adsorption.

### Mitochondria fractionation, Membrane potential and Cellular calcium assay

Mitochondria Isolation Kit, human from Miltenyi Biotec (130-094-532) was used for mitochondria fractionation following the manufacturer’s instructions. For the mitochondrial membrane potential assay, the Mitochondrial Membrane Potential Kit (Sigma Aldrich, MAK159) was used following the manufacturer’s instructions. For cellular calcium assay, Fluo-8 Calcium Assay Kit – No Wash (Abcam, ab112129) was used following the manufacturer’s instructions.

### In-vitro cleavage assay

Pierce^TM^ HRV 3C Protease Solution kit (ThermoFisher, 88946) was used for protein in-vitro cleavage assay following the manufacturer’s instructions. EV-D68 3C-protease (see next section) was used in place of HRV-3C protease, following the same protocol. In brief, A549 cell lysates were incubated with and without recombinant EV-D68 protease at 4°C overnight. The lysates were then prepared for a western blot against MFN-2.

### EV-D68 3C Protein Expression and Purification

The gene corresponding to EV68 3CPro (residues, 1-183) was subcloned into pET28 expression vector (pET28-EV68 3CPro1-183-Hisx6) and was transformed into BL21 (DE3) strain of E. coli competent cells. The cell cultures were grown at 37 °C until OD600 reached 0.6 and induced with 0.5 mM isopropyl ß-d-1-thiogalactopyranoside (IPTG) at 37 °C for 3 hours. Cells were harvested by centrifugation at 4,000 RPM for 20 min at 4°C and collected cell pellets were resuspended in lysis buffer (20 mM Tris/HCl pH 8.0, 300mM NaCl, 5 mM b-mercaptoethanol (BME), 100uL of 5 mg/mL DNase, 10 mM CaCl2, 10 mM MgCl2, 1% glycerol, and 1% B-PERTM), incubated with stirring for 1 hour and lysed by sonication using 50% power (5 mins cycles; 5 secs on, 5 secs off). Cell debris and insoluble proteins were centrifuged at 15,000 RPM for 45 min at 4°C and supernatant was filtered previous to chromatographic purification. EV-D68 3CPro1-183-Hisx6 purification was performed in two steps. First, the filtered lysate was loaded onto immobilized metal affinity chromatography (HiPrepTM IMAC FF 16/10) using buffer A (20mM Tris/HCl pH 8.0, 300 mM NaCl, 5mM BME). EV-D68 3CPro1-183-Hisx6 eluted fractions (120mM - 195mM imidazole) and concentrated for size exclusion chromatography, HiLoad ® 16/600 Superdex200, previously equilibrated with 20 mM HEPES pH 7.5, 150mM NaCl buffer. Eluted fractions (96 to 116 mL) were analyzed by SDS-PAGE, concentrated to 1 mg/mL and stored at −80°C.

### Statistical analysis

GraphPad prism was used for all statistical analysis. Values represent the mean ± standard error of the mean (SEM) of at least 2 to 3 independent repeats. Student t-test was used for comparison and a p-value of < 0.05. was considered statistically significant.

### Open Data Repository

The raw data for this manuscript are available at the Open Science Framework (OSF) at the following private link: https://osf.io/43v68/?view_only=07f708f291934fe891d2102166084bcd

This link is for review only and should not be shared. Upon publication, data will be publicly available at OSF.

## Acknowledgements.

We thank the members of the Jackson, Frieman, and Coughlan labs for weekly discussion sessions. This work was funded by NIH/NIAID grants R01AI141359 and R01AI104928 to W.T.J. N.P. is supported by NIH/NIAID training grant AI095190.

---

**Supplemental Figure 1.**
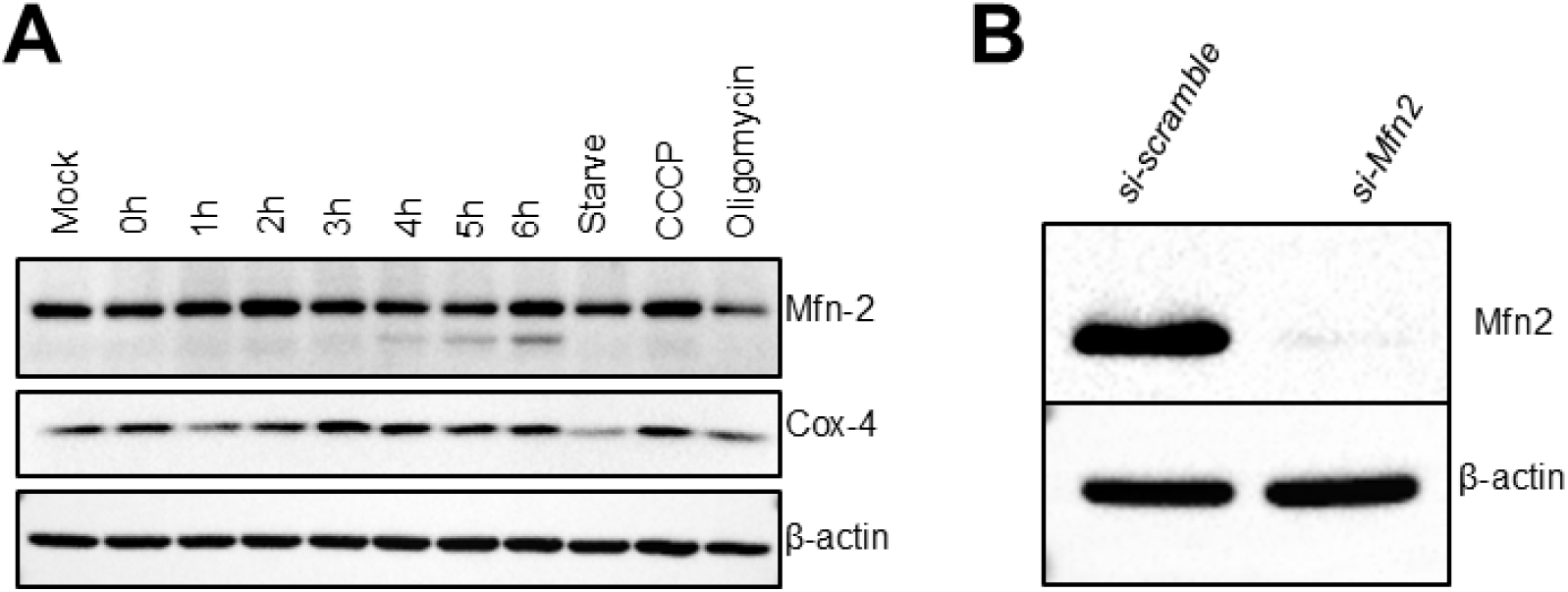
EV-D68 cleaves mitofusin-2 in A549 cells.

